# Gene co-expression modules underlying polymorphic and monomorphic zooids in the colonial hydrozoan, *Hydractinia symbiolongicarpus*

**DOI:** 10.1101/006072

**Authors:** David C. Plachetzki, M. Sabrina Pankey, Brian R. Johnson, Eric J. Ronne, Artyom Kopp, Richard K. Grosberg

## Abstract

Advances in sequencing technology have forced a quantitative revolution in Evolutionary Biology. One important feature of this renaissance is that comprehensive genomic resources can be obtained quickly for almost any taxon, thus speeding the development of new model organisms. Here we analyze 20 RNA-seq libraries from morphologically, sexually, and genetically distinct polyp types from the gonochoristic colonial hydrozoan, *Hydractinia symbiolongicarpus* (Cnidaria). Analyses of these data using Weighted Gene Co-expression Networks highlights deeply conserved genetic elements of animal spermatogenesis and demonstrate the utility of these methods in identifying modules of genes that correlate with different zooid types across various statistical contrasts. RNAseq data and analytical scripts described here are deposited in publicly available databases.

## Introduction

Next Generation Sequencing (NGS) is revolutionizing organismal biology and holds particular promise for understanding the genetic mechanisms that give rise to complex morphological traits (Meng, et al. 2013; Reed et al 2011; Siebert, et al. 2011). One common application of NGS is RNAseq. In RNAseq, sequencing libraries are prepared from mRNAs extracted from tissues of interest and sequenced on a massively parallel platform such as Illumina (Mortazavi, et al. 2008). By mapping individual sequencing reads back to genomic or transcriptome assemblies, quantitative estimates of the expression levels of individual genes can be assessed statistically (Wang, et al. 2009). Comparisons of differential gene expression between tissues of interest are now ubiquitous in biological research, but additional information exists in these large and complex datasets. For instance, genes that underlie a specific biological function are often expressed in a correlated fashion (Miller, et al. 2008; Stuart, et al. 2003). Thus, analytical procedures that focus on identifying such correlations such as Weighted Gene Co-expression Network Analysis (WGCNA; (Langfelder and Horvath 2008), are a powerful means for identifying the key genetic modules underling the trait of interest. Here we test the utility of the WGCNA (Langfelder and Horvath 2008) in characterizing the genetic modules that underlie specific morphological phenotypes (e.g. zooid types) using an extensive RNAseq dataset derived from the modular colonial hydrozoan, *Hydractinia symbiolongicarpus*.

*H. symbiolongicarpus* is a powerful model system for integrative studies in organismal biology for several reasons. First, it is amenable to laboratory husbandry and the full reproductive cycle can be accomplished in the laboratory within roughly three months - a shorter period from generation to generation than other popular model taxa like mice (Silver 1995) or zebrafish (Detrich, et al. 2011). Second, colonies are easily crossed, facilitating the sorts of genetics studies that are difficult in other hydrozoan models such as *Hydra* species. Further, *H. symbiolongicarpus*, like other colonial invertebrates, shows a remarkable capacity for regeneration, allowing individual colonies to be sub-cloned by surgical fragmentation and stocked in perpetuity. In addition, the recent development of transgenic approaches in the congener, *Hydractinia echinata* (Kunzel, et al. 2010), opens the possibility of functional genomics. Finally, transcriptome data for *H. echinata* are available (Soza-Ried, et al. 2010) and a draft genome sequence of *H. symbiolongicarpus* is in preparation (Schnitzler, et al. 2014).

Colonies of *H. symbiolongicarpus* consist of up to four distinct types of polyps, including feeding gastrozooids, reproductive gonozooids, defensive dactylozooids, and sensory spiral zooids, all of which develop from basal ectodermal mat or network of stolons (Frank, et al. 2001; McFadden, et al. 1984). Here we characterize an extensive RNAseq dataset comprising 20 sequencing libraries representing multiple replicates of male and female gastrozooids and gonozooids from different genotypes. We describe a workflow for the analysis of these data using the WGCNA statistical package (Langfelder and Horvath 2008) and show that sufficient signal exists in our RNAseq dataset to construct networks of correlated gene expression. Many of the genetic modules recovered by these methods correlate with specific zooid morphologies. Our analyses provide the foundation for extending our understanding of the modules of functionally related genes that underlie specific morphological features, such as the different types of zooids in *H. symbiolongicarpus* and other modular organisms that exhibit morphological polymorphism at the level of zooids or polyps.

Raw data furnished by this study can be accessed at Sequence Read Archive (http://www.ncbi.nlm.nih.gov/sra) (Bioproject PRJNA244078). Data matrices and scripts for statistical analyses are available at https://bitbucket.org/plachetzki/plachetzki-et-al.-sicb-2014, and sequence files associated with this analysis are located at dryad (doi:10.5061/dryad.98pt3).

## Methods

### Animal Stocks and Husbandry

We collected colonies of *H. symbiolongicarpus* inhabiting shells of the intertidal hermit crab, *Pagurus longicarpus*, from Barnstable Harbor, MA in July of 2010. Animal husbandry was conducted as previously described (Grosberg, et al. 1996). Briefly, we maintained the hermit crabs and their associated *H. symbiolongiocarpus* in a 200-gallon recirculating seawater system (held at 18°C) at the University of California, Davis. We fed the carbs and their associated colonies daily to repletion with 2-3 day old cultures of brine shrimp (*Artemia salina*). We mated single pairs of male and female colonies by isolating crabs carrying single, sexually mature male or female colonies in the dark for 24 hours, then placing them together in two-liter containers of filtered (0.44 µm) seawater (FSW) at room temperature (RT) under bright light for 4 hours. We collected ≈ 500 fertilized embryos from each mating, and transferred them into clean seawater in sterile petri dishes. We transferred the developing embryos daily into clean FSW. After 55 hours we induced metamorphosis by treating planulae with a solution of 55 mM CsCl in FSW (Muller 1973). We transferred metamorphosing planulae to borosilicate glass plates submerged in FSW, where they continued to develop into primary feeding polyps. We fed the resulting colonies daily to repletion with 2-3 day old *Artemia* until the colonies reached sexually maturity after five to eight weeks. We selected two male and two female full-sibling offspring for extractions of RNA.

### Extraction of RNA, Construction of RNAseq Libraries, and Sequencing

We used fine scissors to excise gonozooids and gastrozooids from living colonies, and placed the polyps directly into Trizol (Invitrogen) on ice. We immediately extracted RNA after removal, following manufacturers protocols (Invitrogen). We extracted three replicate total-RNA samples from each type of polyp (male gastrozooid, male gonozooid, female gastrozooid, female gonozooid) from four full-sibling individuals (two male and two female colonies, each), for a total of 24 extractions. We eluted total RNA and stored the RNA at -80° C in RNAse-free H_2_O. We assessed RNA quality using a Bioanalyzer and the RNA 1000 kit (Agilent). We prepared RNAseq libraries using the TrueSeq kit (version 2, Illumina) with 200 – 500 ng of total RNA per library, following the manufacturer’s protocols. We used a Bioanalyzer and DNA 100 chips (Agilent) to determine physical quality of sequencing libraries. Sequencing and de-multiplexing were carried out at the University of California, Davis Genome Center on two lanes of HiSeq 2000 (paired end 100) with 12 RNAseq libraries per lane. Four of our RNAseq datasets had low read content, and we eliminated them from subsequent analysis (see Table 1).

**Table 1.** Description of RNAseq datasets from *Hydractinia symbiolongicarpus* used in the current study. Sequences deposited under Bioproject PRJNA244078.

| **RNAseq Dataset** | **Sex** | **Zooid Type** | **Genotype** | **SRA accession** |
| --- | --- | --- | --- | --- |
| DP1 | Male | Gastrozooid | 48 | SRS598802 |
| DP2 | Male | Gonozooid | 5 | SRS598818 |
| DP3 | Female | Gastrozooid | 17 | SRS599147 |
| DP4 | Female | Gonozooid | 17 | SRS599148 |
| DP5 | Male | Gastrozooid | 48 | SRS599223 |
| DP6 | Male | Gonozooid | 48 | SRS599224 |
| DP7 | Female | Gastrozooid | 73 | SRS599225 |
| DP8 | Female | Gonozooid | 73 | SRS599227 |
| DP9 | Male | Gastrozooid | 5 | SRS599283 |
| DP10 | Male | Gonozooid | 5 | SRS599284 |
| DP11 | Female | Gonozooid | 17 | SRS599285 |
| DP12 | Male | Gastrozooid | 48 | SRS599287 |
| DP13 | Male | Gastrozooid | 73 | SRS599872 |
| DP15 | Female | Gonozooid | 73 | SRS599873 |
| DP16 | Male | Gastrozooid | 5 | SRS599874 |
| DP17 | Male | Gonozooid | 5 | SRS599875 |
| DP18 | Female | Gastrozooid | 17 | SRS599876 |
| DP19 | Female | Gonozooid | 17 | SRS601014 |
| DP20 | Male | Gonozooid | 48 | SRS601015 |
| DP21 | Female | Gastrozooid | 17 | SRS601017 |

### Bioinformatics: Filtering and Assembly of Raw Data

We first cleaned low-quality reads from the raw sequence data for each RNAseq dataset using fastq_quality_filter version 0.013 and the first 10 base pairs of each read were removed using fastx_trimmer version 0.013 (http://hannonlab.cshl.edu/fastx_toolkit/index.html). Sequencing adaptors were removed using cutadapt (Martin 2011). Comparisons between pre-cleaning and post-cleaning datasets were made using fastqc version 0.10.0 (http://www.bioinformatics.babraham.ac.uk/projects/fastqc/). Following cleaning, we used Trinity version 2013.11.10 (Grabherr, et al. 2011) under default parameters to assemble non-redundant datasets with the highest number of reads. We then mapped reads from individual, cleaned RNAseq datasets to contigs using bowtie version 1.0.1 (Langmead, et al. 2009) within RSEM version 1.1.15 (Li and Dewey 2011). Mapping efficiency was assessed using the flagstat command in samtools version 0.1.19 (Li, et al. 2009). The raw transcriptome assembly and the resulting matrix of un-normalized expected count data are available at the dryad link associated with this work.

### Bioinformatics: Statistical Analytics

We analyzed the counts matrix from all of 20 datasets using Weighted Gene Co-expression Network Analysis (WGCNA) version 1.36 (Langfelder and Horvath 2008). First, we discarded all contigs that included fewer than 10 counts in any library. The resulting matrix included 3,539 contigs. Next, we normalized count data using the model based calcNormFactors function in EdgeR version 3.6.2 (Robinson, et al. 2010). Hierarchical clustering was then used to assess similarity between the resulting 20 RNAseq datasets using flashClust in R version 3.0.0 (R_Development_Core_Team 2008). As none of our datasets exceeded a threshold of 0.9% divergence in Euclidian distance, none were considered outliers and all were included in the following analyses. Soft thresholding analyses of powers and WGCNA analyses were conducted as per the authors’ instructions using the pickSoftThreshold function in WGCNA. Once modules of co-expressed genes were estimated, statistical analyses included: i) tests of association between individual genes and proposed module membership, ii) tests of association between individual genes and sex or zooid type, and iii) analyses of variance (ANOVA) to test for the effect of zooid type and sex on expression of each of the elucidated modules. P-values were corrected for multiple tests using a false discovery rate of 0.05. Finally, the sequences of individual modules of co-expressed genes that resulted from this analysis were BLASTed against the refseq_protein database (Altschul, et al. 2009) with an expect value of 1.0 × 10^−3^ and annotated using BLAST2GO (Conesa, et al. 2005).

## Results

### RNAseq Data

Twenty *Hydractinia symbiolongicarpus* RNAseq datasets (paired end 100bp, HiSeq 2000) were prepared during this study. These datasets comprise multiple replicates of male and female gastrozooids and gonozooids, with two genotypes per sex (genotypes 5, 48, 17 and 73). Table 1 describes these data and their publicly available accessions, which together comprise > 2.5 × 10^8^ reads.

Assembly in Trinity produced 120,196 contigs (transcripts). Twenty-five per cent of the total sequence length from the assembly is contained in 8,519 contigs of 1,556 bp or greater, 50% of total sequence length from the assembly is contained in 24,514 contigs of 890 bp or greater, and 75% of total sequence length from the assembly is contained in 54,696 contigs of 432 bp or greater. On average 88% of read data across 20 cleaned RNAseq datasets mapped to the assembly.

Figure 1 shows a dendrogram of these 20 RNAseq libraries, following the removal of contigs that had fewer than 10 counts per contig for each library. Each type of polyp, including female gonozooids, male gonozooids, and gastrozooids were recovered each as unique clusters, the gastrozooid cluster being comprised of both sexes. The genotype of a given RNAseq library was not a strong determinant in its clustering, as would be expected if genotype played a limited role in driving differences in gene expression between types of polyps.

**Figure 1.**
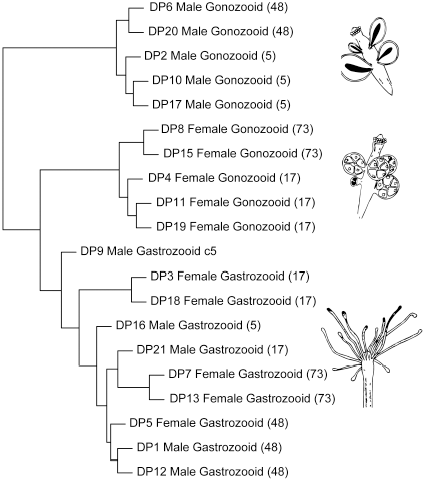
Cluster dendrogram of the 20 RNAseq datasets from *Hydractinia symbiolongicarpus* described in this paper. The branching pattern shown here is drawn from overall Euclidean distances in gene expression between each library after pruning and normalization of data (see methods). This analysis clusters male and female gonozooids into separate groups, while gastrozooids of both sexes are clustered into a single group. Dendrogram based on 3,549 transcripts that had greater than 10 reads per contig in each library.

### WGCNA

The first step in weighted co-expression gene network analysis (WGCNA) is the selection of β, the soft thresholding power used to calculate the adjacency matrix, or co-expression network, of genes. Using the automated approach to network construction (Langfelder and Horvath 2008; Zhang and Horvath 2005) we selected a power of 9 for β, which is the inflection point in proposed power values where the scale-free topology model fit to our data begins to stop improving with increases in power (Fig. 2). Using this value for β, our data were partitioned into 10 gene co-expression modules that included between 38 and 1258 transcripts (Fig 3A). Fifteen transcripts, out of the original 3,549 did not fall into any significant module of gene co-expression.

**Figure 2.**
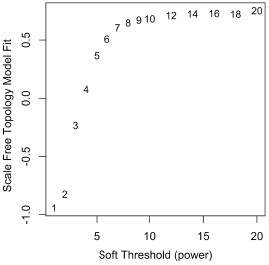
Selection of soft thresholding power β. Our filtered dataset was fit to a scale free model (x axis) of proposed values for β ranging from 1–20. We selected a value of 9 for β as it reflects the inflection point where model fit begins to decrease with increases in power.

**Figure 3.**
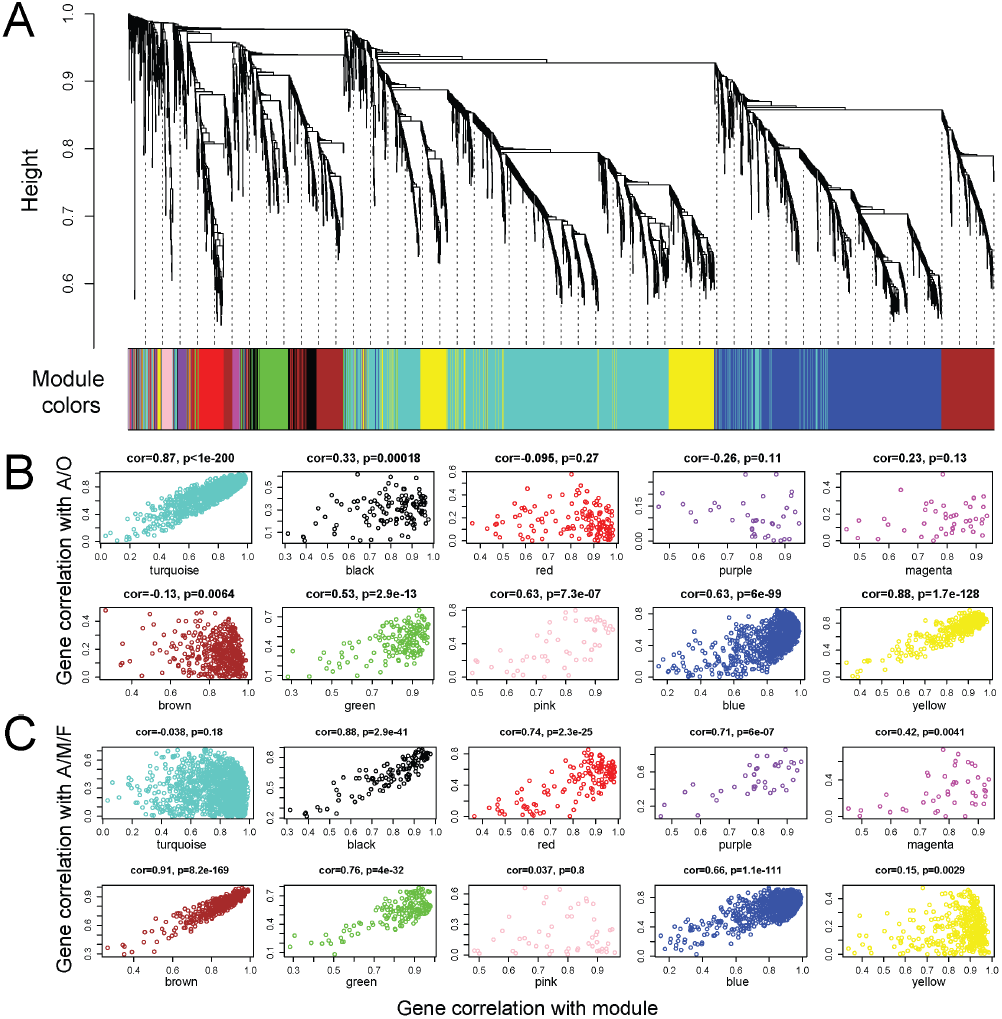
Cluster dendrogram and trait-correlations of 3,549 transcripts analyzed using WGCNA (Langfelder and Horvath 2008). **A**, Cluster dendrogram showing the relationships of all transcripts and their assignment to specific modules (colored bars) of correlated gene expression. **B**, Scatterplots showing the relationship between trait-gene correlation and module-gene correlations under the A/O trait scheme (gastrozooid (A) and gonozooid (O)). Here the eigenvectors estimated in WGCNA for each gene are first correlated with a given gene’s module of membership (x axis, denoted by bars of different colors in **A**) and then those values are correlated with a two-state conditional variable A/O (y axis). Modules with high-scoring correlations are those in which within-module variation is explained by the A/O contrast. **C**, Scatterplots showing the relationship between trait-gene correlation and module-gene correlations under the A/MO/FO trait scheme (gastrozooid (A), male gonozooid (MO), and female (FO)). As in **B**, the eigenvectors estimated in WGCNA for each gene are first correlated with a given gene’s module of membership (x axis) and then those values are correlated with the three-state conditional variable A/MO/FO (y axis). Modules with high-scoring correlations are those in which within-module variation is explained by the A/MO/FO contrast.

Genes included in specific modules may share similar biological functions. Thus, understanding the composition of specific modules may inform our understanding of the differences in gene expression that underlie the different types of zooids in *H. symbiolongicarpus*. In order to assign modules of co-expressed transcripts to specific traits (i.e. gastrozooids and gonozooids of different sexes), we first verified a gene’s module membership by testing the correlation between the individual gene’s expression and the overall expression eigenscore of its assigned module. We then tested the association between module eigenscores and trait (male gonozoid (MO), female gonozooid (FO) or gastrozooid (A)). Figure 3B shows the results of these correlations when the zooid type A/O (gastrozooid A and gonozooid O) is used as a correlate of membership in a gene module. This categorization scheme disregards sex differences between zooid samples. These analyses suggest that the turquoise, pink, blue, and yellow modules are strongly associated with A/O zooid types and therefore may represent genetic modules underlying some of the morphological differences in these zooid types (Fig. 3B). However, other modules that do not yield high correlations between module membership and A/O trait-distinctions may be explained by other possible trait categorization schemes. For instance, genes included in the red, brown, and green modules show a low correlation to zooid type under the A/O scheme, but a high correlation to zooid type once zooids are categorized as either gastrozooid, male gonozooid, or female gonozooid (A/MO/FO) (Fig. 3C).

In order to test whether zooid type and/or sex has significant explanatory power to predict the expression of our estimated genetic modules, we conducted an analysis of variance on each module (Fig. 4). These analyses suggest that the turquoise module is significantly associated with gastrozooid type (zooid: *F* = *100*, *p* < *0.001*); the magenta and brown modules with sex differences (magenta: *F* = *6.5*, *p* = *0.0424*; brown: *F* = *155*, *p* < *0.001*); the green module is strongly associated with the female gonozooid type (sex: *F* = *35.4*, *p* = *0.0001*; zooid: *F* = *30.1*, *p* = *0.0002*); and the blue module associates strongly with the male gonozooid type (sex: *F* = *12.5*, *p* = *0.0069*; zooid: *F* = *16.1*, *p* = *0.0034*). Support for this association between the between blue module and male gonozooid polyp type comes from GO analyses of the blue module, which reveal a number of genes associated with spermatogenesis and meiosis (Table 2). The sequences of the transcripts from each module are given at the dryad link associated with this publication; however, a detailed global analysis of all the genes contained in each module will be explicated in a forthcoming publication.

**Figure 4.**
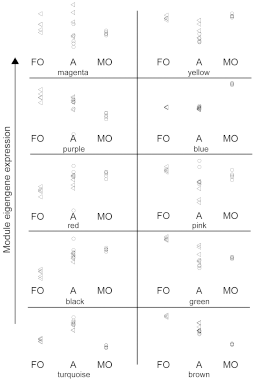
Module eigengene expression distributions across libraries for each zooid type. Eigengene expression values for each of 20 libraries were plotted for each module by zooid type. Our results from ANOVA suggest that the turquoise module strongly corresponds to the gastrozooids of both sexes (A), the brown, and green modules correspond to the female gonozooid (FO) trait, and the blue module corresponds to the male gonozooid (MO) trait. circles = male; triangles = female.

**Table 2.** *Hydractinia symbiolongicarpus* contigs (transcripts) with gene ontological terms related to male-specific traits from the blue, male gonozooid-specific, module of co-expressed genes.

| **Contig Name** | **Closest BLAST Hit** | **Process (P) and Cellular Compartment (C)** | **GO #** |
| --- | --- | --- | --- |
| comp54693_c0 | bardet-biedl syndrome 4 protein | spermatid development | GO:0007286 |
| comp36203_c0 | dihydrolipoyl mitochondrial-like | sperm capacitation | GO:0048240 |
| comp50065_c0 | e3 ubiquitin-protein ligase herc2 | Spermatogenesis | GO:0007283 |
| comp58851_c0 | piwi-like protein 1 | Spermatogenesis | GO:0007283 |
| comp40539_c0 | sorbin and sh3 domain-containing protein 2 isoform x3 | spermatid development | GO:0007286 |
| comp54701_c0 | sperm surface protein sp17 | binding of sperm to zona pellucida | GO:0007339 |
| comp54434_c0 | calreticulin-like | regulation of meiosis | GO:0040020 |
| comp59793_c0 | structural maintenance of chromosomes protein 3 | Meiosis | GO:0007126 |

## Discussion

We describe the data and their deposition in publicly available databases from 20 RNAseq libraries derived from multiple replicates of male and female gastrozooids and gonozooids from the colonial hydrozoan *H. symbiolongicarpus*. Our analyses suggest that the WGCNA approach to uncovering genes associated with specific morphological features may be a powerful tool for organismal biology. We recovered gene modules that correlate strongly with specific types of zooids. In particular, the blue module correlates with the male gonozooid trait and this module contains numerous genes associated with spermatogenesis and meiosis (Table 2).

The modules of correlated gene expression estimated from WGCNA are *a priori* naïve to trait-states and associations between the specific modules identified and trait-states are driven by nuances in the expression data. This renders the WGCNA approach a valuable tool for identifying the key genetic constituents underlying traits where little is known, such as the polymorphic polyp morphologies of *H. symbiolongicarpus*. In addition, modules that show no correlation with any particular state of a trait do not represent failures in the procedure. Such modules may represent molecular functions that are shared across states but have correlated expression nonetheless, such as metabolic pathways.

All comparative transcriptomics analyses are likely to include false positive and negative results. Therefore, analyses such as these represent only a first step in identifying the genes that function to specify different polyp morphologies. Future analyses will investigate the expression patterns and molecular functions of selected candidate genes recovered in the modules of correlated gene expression described here. In addition, our present analyses of gene ontology are based on BLAST (Altschul, et al. 1997) searches of similarity against accessioned databases. While useful, such analyses conflate orthology and parology. Future analyses will apply a more robust phylogenetic approach aimed at disentangling orthology from parology in genes of interest. Only then may we be able to address the evolutionary significance of our findings as they relate to the origins of polymorphic polyp types and gametogenesis.

## Acknowledgements

We are indebted to Brenda B. Cameron for assistance and training in animal husbandry. The manuscript benefited greatly from helpful comments from two anonymous reviewers and the editorial staff at ICB.

## References Cited

Altschul SF, Gertz EM, Agarwala R, Schaffer AA, Yu YK 2009. PSI-BLAST pseudocounts and the minimum description length principle. Nucleic Acids Res 37: 815–824. doi: 10.1093/nar/gkn981

Altschul SF, Madden TL, Schaffer AA, Zhang J, Zhang Z, Miller W, Lipman DJ 1997. Gapped BLAST and PSI-BLAST: a new generation of protein database search programs. Nucleic Acids Res 25: 3389–3402.

Conesa A, Gotz S, Garcia-Gomez JM, Terol J, Talon M, Robles M 2005. Blast2GO: a universal tool for annotation, visualization and analysis in functional genomics research. Bioinformatics 21: 3674–3676. doi: 10.1093/bioinformatics/bti610

Detrich HW, Westerfield M, Zon LI. 2011. The Zebrafish: Genetics, Genomics and Informatics: Academic Press.

Frank U, Leitz T, Muller WA 2001. The hydroid Hydractinia: a versatile, informative cnidarian representative. Bioessays 23: 963–971. doi: 10.1002/bies.1137

Grabherr MG, Haas BJ, Yassour M, Levin JZ, Thompson DA, Amit I, Adiconis X, Fan L, Raychowdhury R, Zeng Q, Chen Z, Mauceli E, Hacohen N, Gnirke A, Rhind N, di Palma F, Birren BW, Nusbaum C, Lindblad-Toh K, Friedman N, Regev A 2011. Full-length transcriptome assembly from RNA-Seq data without a reference genome. Nat Biotechnol 29: 644–652. doi: 10.1038/nbt.1883

Grosberg RK, Levitan DR, Cameron BB 1996. Evolutionary genetics of allorecognition in the colonial hydroid Hydractinia symbiolongicarpus. Evolution 50: 2221–2240.

Kunzel T, Heiermann R, Frank U, Muller W, Tilmann W, Bause M, Nonn A, Helling M, Schwarz RS, Plickert G 2010. Migration and differentiation potential of stem cells in the cnidarian Hydractinia analysed in eGFP-transgenic animals and chimeras. Dev Biol 348: 120–129. doi: 10.1016/j.ydbio.2010.08.017

Langfelder P, Horvath S 2008. WGCNA: an R package for weighted correlation network analysis. BMC Bioinformatics 9: 559. doi: 10.1186/1471-2105-9-559

Langmead B, Trapnell C, Pop M, Salzberg SL 2009. Ultrafast and memory-efficient alignment of short DNA sequences to the human genome. Genome Biol 10: R25. doi: 10.1186/gb-2009-10-3-r25

Li B, Dewey CN 2011. RSEM: accurate transcript quantification from RNA-Seq data with or without a reference genome. BMC Bioinformatics 12: 323. doi: 10.1186/1471-2105-12-323

Li H, Handsaker B, Wysoker A, Fennell T, Ruan J, Homer N, Marth G, Abecasis G, Durbin R, Genome Project Data Processing S 2009. The Sequence Alignment/Map format and SAMtools. Bioinformatics 25: 2078–2079. doi: 10.1093/bioinformatics/btp352

Martin M 2011. Cutadapt removes adapter sequences from high-throughput sequencing reads. EMBnet.journal 17: 10–12.

McFadden CS, McFarland MJ, Buss LW 1984. Biology of Hydractiniid Hydroids. 1. Colony Ontogeny in Hydractinia echinata (Flemming). Biological Bulletin 166: 54–67.

Meng F, Braasch I, Phillips JB, Lin X, Titus T, Zhang C, Postlethwait JH 2013. Evolution of the eye transcriptome under constant darkness in Sinocyclocheilus cavefish. Mol Biol Evol 30: 1527–1543. doi: 10.1093/molbev/mst079

Miller JA, Oldham MC, Geschwind DH 2008. A systems level analysis of transcriptional changes in Alzheimer’s disease and normal aging. J Neurosci 28: 1410–1420. doi: 10.1523/JNEUROSCI.4098-07.2008

Mortazavi A, Williams BA, McCue K, Schaeffer L, Wold B 2008. Mapping and quantifying mammalian transcriptomes by RNA-Seq. Nat Methods 5: 621–628. doi: 10.1038/nmeth.1226

Muller WA 1973. Induction of metamorphosis by bacteria and ions in the planulae of Hydractinia echinata; an approach to the mode of action. PUBLICATIONS OF THE SETO MARINE BIOLOGICAL LABORATORY 20: 195–208.

R Development Core Team. 2008. R: A language and environment for statistical computing. Vienna, Austria.

Reed RD, Papa R, Martin A, Hines HM, Counterman BA, Pardo-Diaz C, Jiggins CD, Chamberlain NL, Kronforst MR, Chen R, Halder G, Nijhout HF, McMillan WO 2011. optix drives the repeated convergent evolution of butterfly wing pattern mimicry. Science 333: 1137–1141. doi: 10.1126/science.1208227

Robinson MD, McCarthy DJ, Smyth GK 2010. edgeR: a Bioconductor package for differential expression analysis of digital gene expression data. Bioinformatics 26: 139–140. doi: 10.1093/bioinformatics/btp616

Schnitzler CE, Sanders SM, Plickert G, Seoighe C, Buss LW, Wollfsburg TG, Nicotra ML, Mullikin JC, Cartwright P, Frank U, Baxevanis AD editors. Society for Integrative and Comparative Biology. 2014 Austin.

Siebert S, Robinson MD, Tintori SC, Goetz F, Helm RR, Smith SA, Shaner N, Haddock SH, Dunn CW 2011. Differential gene expression in the siphonophore Nanomia bijuga (Cnidaria) assessed with multiple next-generation sequencing workflows. PLoS One 6: e22953. doi: 10.1371/journal.pone.0022953

Silver LM. 1995. Mouse genetics: concepts and applications. New York: Oxford University Press.

Soza-Ried J, Hotz-Wagenblatt A, Glatting KH, del Val C, Fellenberg K, Bode HR, Frank U, Hoheisel JD, Frohme M 2010. The transcriptome of the colonial marine hydroid Hydractinia echinata. FEBS J 277: 197–209. doi: 10.1111/j.1742-4658.2009.07474.x

Stuart JM, Segal E, Koller D, Kim SK 2003. A gene-coexpression network for global discovery of conserved genetic modules. Science 302: 249–255. doi: 10.1126/science.1087447

Wang Z, Gerstein M, Snyder M 2009. RNA-Seq: a revolutionary tool for transcriptomics. Nat Rev Genet 10: 57–63. doi: 10.1038/nrg2484

Zhang B, Horvath S 2005. A general framework for weighted gene co-expression network analysis. Stat Appl Genet Mol Biol 4: Article17. doi: 10.2202/1544-6115.1128

